# Structural basis for the inhibition of translation through eIF2α phosphorylation

**DOI:** 10.1101/503979

**Authors:** Yuliya Gordiyenko, José Luis Llácer, Venki Ramakrishnan

**Affiliations:** MRC Laboratory of Molecular Biology, United Kingdom; Instituto de Biomedicina de Valencia del Consejo Superior de Investigaciones Científicas, Valencia, Spain

**Author notes:** these authors contributed equally to this work (Yuliya Gordiyenko, José Luis Llácer). corresponding author: Dr. José Luis Llácer, Instituto de Biomedicina de Valencia (IBV-CSIC), C/ Jaime Roig 11, Valencia-46010, Spain.

## Abstract

One of the responses to stress by eukaryotic cells is the down-regulation of protein synthesis by phosphorylation of translation initiation factor eIF2. Phosphorylation results in low availability of the eIF2 ternary complex (eIF2-GTP-tRNA_i_) by affecting its interaction with its GTP-GDP exchange factor eIF2B. We have determined the cryo-EM structure of eIF2B in complex with phosphorylated eIF2 at an overall resolution of 4.15 Å. Two eIF2 molecules bind opposite sides of an eIF2B hetero-decamer through eIF2α-D1, which contains the phosphorylated Ser51. eIF2α-D1 is mainly inserted between the N-terminal helix bundle domains of δ and α subunits of eIF2B. Phosphorylation of Ser51 enhances binding to eIF2B through direct interactions of phosphate groups with residues in eIF2Bα and indirectly by inducing contacts of eIF2α helix 58-63 with eIF2Bδ leading to a competition with Met-tRNA_i_.

## Introduction

In eukaryotes initiation of protein synthesis is tightly regulated by a number of translation initiation factors (eIFs) including the GTPase eIF2. During initiation, the GTP-bound eIF2 forms a ternary complex (TC) with Met-tRNA_i_^Met^, and together with other initiation factors binds the 40S ribosomal subunit, forming the 43S pre-initiation complex (PIC). Another initiation factor is the GTPase activating protein (GAP) eIF5, which promotes GTP hydrolysis by eIF2 ^1,2,3,4^, helps to locate the AUG start codon at the P site during scanning along mRNA ^5^ After the PIC recognition of the initiation codon, inorganic phosphate is released ^3^ and the GDP-bound eIF2 dissociates from the 40S along with most other initiation factors, and the subsequent binding of eIF5B promotes joining of the 60S and the start of the protein synthesis. For multiple rounds of initiation to occur, the GDP on eIF2 has to be exchanged for GTP. This reaction is catalysed by a guanine nucleotide exchange factor (GEF) eIF2B.

eIF2 is a protein complex, comprising three subunits, eIF2α, eIF2β and eIF2γ. Of these, eIF2γ has the catalytic site for GTPase activity and binds and recognises the acylated acceptor arm of the Met-tRNA_i_ ^Met 6,7^ eIF2β forms part of the nucleotide binding pocket in eukaryotes ^7^ and eIF2α is inserted in the E site of the 40S subunit during translation initiation while being bound to Met-tRNA_i_^Met 7,8,9^ and also has regulatory function ^10, 11^. In response to various stress conditions eukaryotic cells regulate protein synthesis by phosphorylation of serine 51 on the eIF2α, thereby converting eIF2 from a substrate to an inhibitor of its GEF, eIF2B ^12, 13^. This highly conserved mechanism, called integrated stress response (ISR) in mammals or general amino acid control (GAAC) in yeast, shuts down bulk protein synthesis ^10 14^ due to the low availability of the TC, and redirects cell resources to adaptive and survival pathways ^15,16, 17, 18^. Deregulation of eIF2B function in humans leads to hypomyelination and neurodegenerative disorders ^19, 20^.

The mechanism of nucleotide exchange by eIF2B and its inhibition by eIF2α phosphorylation has been a matter of considerable debate^12, 21–25,26, 27^. The regulatory subunits α, β, δ are homologous with a similar fold and form the hexameric core of eIF2B, while the catalytic subunits γ and ε assemble into heterodimers and bind peripherally on two opposite sides of the regulatory hexamer as shown in the x-ray structure of *S. pombe* eIF2B ^28^ and cryoEM structures of human eIF2B ^26, 27^. eIF2B γ and ε are homologous to each other and have two domains in common - a pyrophosphorylase like domain (PLD) and a left-handed β helix (LβH) domain ^29^. eIF2Bε in addition has a C-terminal heat domain extension ^30^ -ε-cat, which itself possesses catalytic activity ^31^. This structural complexity makes it more difficult to understand the mechanism of action and regulation of eIF2B.

The interactions of eIF2 with eIF2B have been extensively investigated biochemically and genetically by mutagenesis of both factors ^32, 33, 34, 35, 36, 37^. In addition, the thermodynamics of eIF2-GDP recycling to the TC has also been studied ^24^. Nevertheless, in the absence of a structure of the eIF2B -eIF2 complex, details of the mechanism of nucleotide exchange and its inhibition by eIF2α phosphorylation remain unclear.

Here we have determined a cryoEM structure of eIF2B in complex with the GDP-bound form of eIF2 phosphorylated at Ser51 on the α subunit, which sheds light on the molecular interactions between the two molecules and provides a basis for understanding the regulation of translation by eIF2α phosphorylation.

## Results

### An overall structure of eIF2B-eIF2(αP) complex

The structure of eIF2B-eIF2(αP) complex was determined to an overall resolution of 4.15 Å at best, although parts of eIF2 show a high degree of conformational heterogeneity. To deal with the conformational heterogeneity, we applied a two-fold C2 symmetry during EM data processing with an averaged position for the eIF2 molecules, and obtained two maps (maps 1 and 2, Fig. S1). Additionally, we obtained four density maps (maps A to D in Fig. S1) without applying any internal symmetry, but at the expense of the overall resolution.

The structure consists of two eIF2 molecules bound to opposite sides of the eIF2B hetero-decamer (Fig. 1a and 1b). Thus the contacts of the two molecules of eIF2 with the eIF2B hetero-decamer are spatially well separated. As judged by the local resolution (Fig. S2), the most stable contact consists of eIF2α domain D1 inserted between the C-terminal helix bundle domains of α and δ regulatory subunits of eIF2B. This interaction is possibly further enhanced by phosphorylation of eIF2α in our complex. Another contact is formed by eIF2γ and eIF2β interacting with the catalytic eIF2B subunits γ and ε (Fig. 1b). This contact has lower local resolution, suggesting that the region has conformational heterogeneity and the interaction is very dynamic.

**Figure 1.**
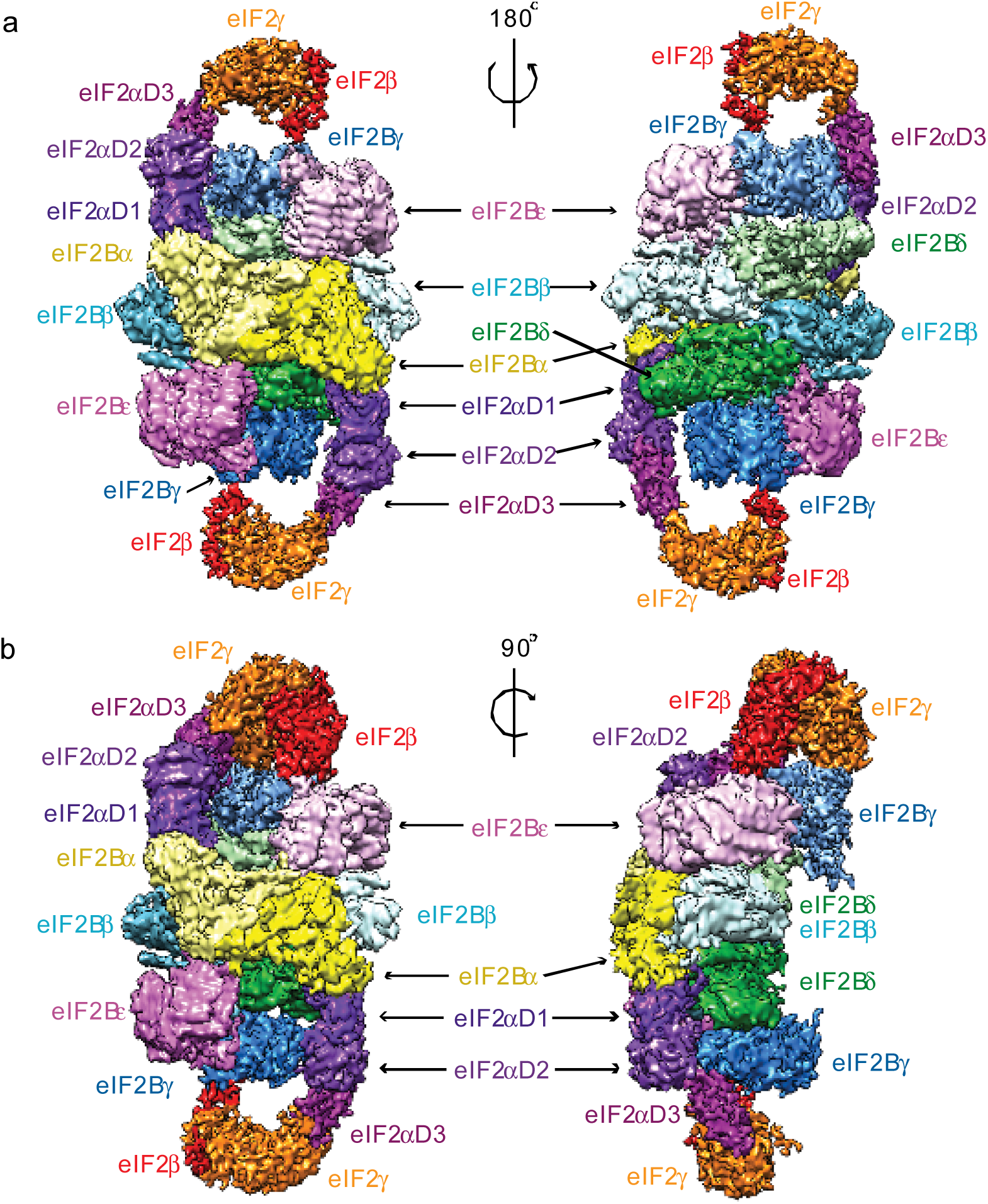
Overview of Cryo-EM structure of eIF2B-eIF2(αP) complex. (a) Two views of the overall cryo-EM map 2 of eIF2B-eIF2(αP) complex at 4.25 Å resolution with different subunits of the complex colour-coded. (b) Two views of the cryo-EM map A of eIF2B-eIF2(αP) complex at 4.6 Å containing clear density for eIF2β subunit on one side of the complex at the top.

In a low-resolution filtered map 2 contoured at lower threshold we could see weak densities around eIF2γ, which cannot be attributed to this subunit (Fig. S1, blue and red masks). Masked classification ^38^ around these densities and eIF2γ allowed us to separate different conformations that eIF2 γ and β adopt in these four different maps. In two of these maps additional low-resolution density could be attributed to the ε-cat heat domain of eIF2Bε (Fig. S1, extra density in map D), but a precise positioning of ε-cat was not possible at this resolution.

### Interaction of the phosphorylated Ser51 on eIF2 with elements of eIF2B

The phosphorylated Ser51 is part of the domain eIF2α-D1, and the structure provides a rationale for why phosphorylation of this residue should inhibit eIF2B function. The domain is inserted between the N-terminal helix bundle domains of δ and α subunits of one set of eIF2B subunits (Fig. 2a and 2b) rather than binding the central cleft of eIF2B as proposed in a previous model ^28^. Interestingly, the cross-links of eIF2α to eIF2Bα and δ obtained for the model ^28^ are in perfect agreement with the binding of eIF2α-D1 in our structure (Fig. 2b), whereas cross-links to eIF2β cannot be explained in the context of our structure. Instead, in agreement with previously identified mutations I118T and S119P in eIF2Bβ that were shown to reduce the effect of eIF2α phosphorylation ^39^, the loop 113–120 of eIF2Bβ (coloured brown), from what could be considered another set of eIF2B subunits, participate in the contact with eIF2α-D1 (Fig. 2b).

**Figure 2.**
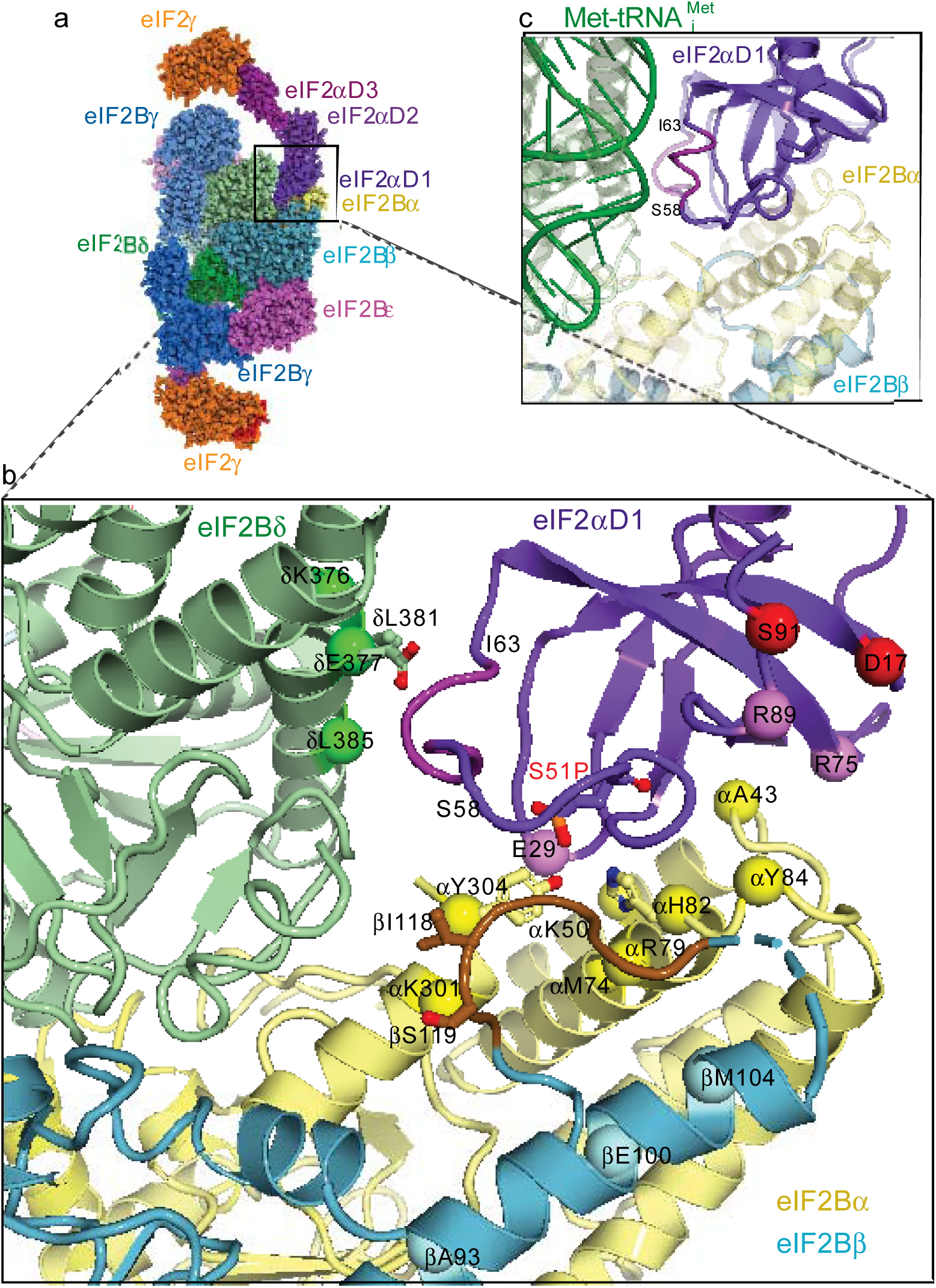
Contacts of eIF2α with the regulatory eIF2B subunits. (a) Model of eIF2B-eIF2(αP) complex fitted in maps 1 and 2. (b) Contacts of eIF2α-D1 with α, β and δ regulatory subunits of eIF2B. Possible residues in contacts with eIF2α S51 phosphate (red sticks) are H82 and Y304 in eIF2Bα (shown in yellow sticks). eIF2Bδ E377 (green sticks) in contact with the 56–63 helix (magenta) of eIF2α affected by the phosphate is also shown. eIF2Bδ E377K overcomes the effect of S51 phosphorylation and eIF2Bβ I118T and S119P (brown sticks) reduce the effect of phosphorylation. Also shown are residues in *S. cerevisiae* eIF2B regulatory subunits α, β and δ corresponding to *S. pombe* residues which cross-linked to eIF2α and residues in eIF2α which cross-linked to eIF2Bα (pink spheres) and to eIF2Bβ (red spheres)^28^. (c) Superposition of eIF2α-D1 in eIF2B-eIF2(αP) complex and in the TC (PDB 3JAP), showing that the same helix 58–63 (coloured magenta) in eIF2α-D1 interacts with both eIF2Bδ and Met-tRNA_i_^Met^ suggesting direct competition for eIF2α-D1.

When compared to the crystal structure of *S. pombe* eIF2B alone ^28^ or cryoEM structures of human ISRIB bound eIF2B ^26, 27^ (Fig. S3a), the binding of eIF2α-D1 in our complex leads to a closure of eIF2B δ and α helix bundle CTD domains around it (Fig. S3b). Closure of the domains also leads to a visible displacement of eIF2Bγ PLD about 5–6 Å outwards (Fig. S3c), making the eIF2B hetero-decamer in the complex with eIF2(αP) elongated by ~ 10–12 Å compared to an apo form ^28^ or ISRIB bound human eIF2B ^26, 27^ (Fig. S3a). The most extensive interaction surface area (844 Å^2^) is between the eIF2α-D1 and eIF2Bα subunits, which would explain why eIF2B mutants lacking an α subunit are not sensitive to eIF2α phosphorylation, as the major part of the binding surface with eIF2α-D1 would be lost.

The density in eIF2α-D1 leading to and including the phosphorylated Ser51 is visible (Fig. 2b and S4), however the arginine-rich loop following this serine seems to be only partially ordered. At this resolution, we cannot establish with complete confidence the interaction partners of Ser51-P because the densities for the side chains around the residue are not absolutely clear. However, the closest residues to the phosphate on Ser51 appear to be eIF2Bα H82 and Y304 (Fig. 2b). Furthermore, in this position the phosphate may affect the conformation of the short α-helix 58–63 after the Arg-rich loop that in turn makes contacts with the eIF2Bδ CTD in our structure (Fig. 2b). eIF2Bδ residues E377 and L381 are likely to be involved in this interaction as mutations E377K and L381Q were shown to overcome the effect of Ser-51 phosphorylation ^33^, suggesting that described mutations would disrupt or weaken this interaction. Indeed, mutation of the residue analogous to E377 in *S. pombe* (D248K) abrogated strong interaction of eIF2(α)P with eIF2B and alleviated inhibition of nucleotide exchange ^28^.

eIF2α phosphorylation is known to increase its binding affinity to eIF2B ^40^, and our structure suggests that this is due to a combination of direct contact of Ser51-P with residues in eIF2Bα (H82 and/or Y304) as well as tighter interaction of the 58–63 α-helix with eIF2Bδ. Interestingly the same helix 58–63 contacts Met-tRNA_i_^Met^ in the TC structure ^7,9,41^, although in the TC, this helix adopts a slightly different conformation (Fig. 2c). This suggests that initiator tRNA and eIF2B may compete for the same binding site on eIF2α, and the altered conformation of the helix upon Ser51 phosphorylation may inhibit the binding of initiator tRNA and displacement and dissociation of eIF2B.

### eIF2 γ and β interactions with catalytic eIF2B subunits

While eIF2α-D1 containing the phosphorylated Ser51 is relatively constrained through its interaction with eIF2B, the domains eIF2 γ and β in the proximity of the catalytic portion of eIF2B have relatively high conformational heterogeneity presumably arising from high mobility (Fig. 1b, 3c and 3d) and do not adopt the same conformation in two eIF2 molecules bound on either side of eIF2B (Fig. 3a). Because of this heterogeneity, which resulted in lower resolution, we cannot be sure whether the GDP that was present in our preparations has been displaced from eIF2γ.

**Figure 3.**
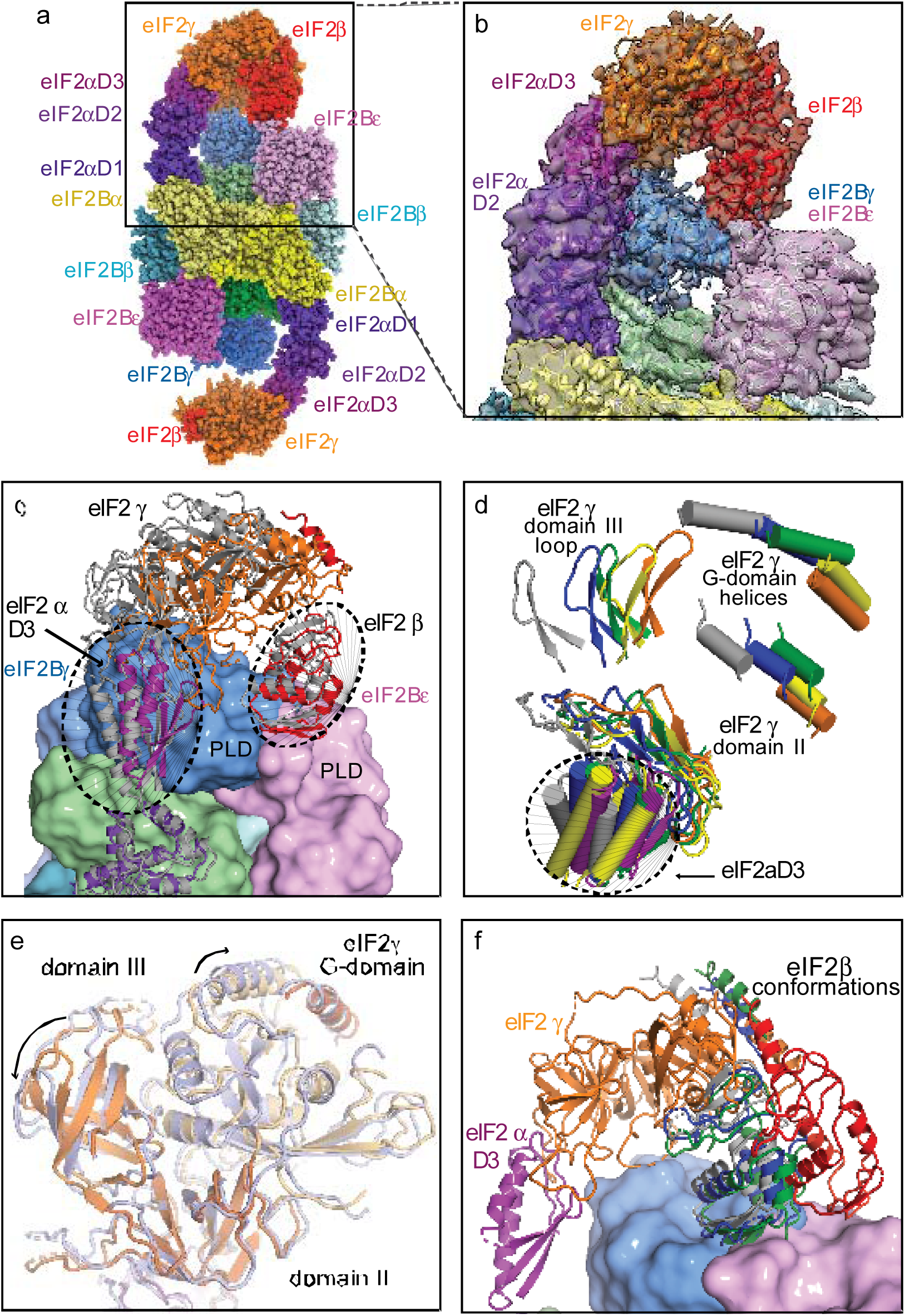
Contacts of eIF2 β and γ with the catalytic subunits of eIF2B. (a) eIF2B-eIF2(αP) complex model in spheres representation fitted in map A showing tilted conformation of eIF2γ which is stabilised by its contact with eIF2Bγ and extended conformation of eIF2β contacting the interface area of the two γ and ε catalytic subunits of eIF2B. (b) Close up view of the model fitting into the density of map A. (c) Modelled positions of eIF2 γ and β subunits after classification, showing extensive movements of these subunits around eIF2B ε and γ PLD domains. For clarity only two eIF2 models are shown (corresponding to maps A - coloured and C - grey). (d) Modelled positions of eIF2γ subunit and eIF2α-D3 in all three maps (map 1-orange for eIF2γ and purple for eIF2α-D3, map A- blue, map B - green, map C - grey, map D - yellow). For clarity only few elements in each of the eIF2γ domains are shown. (e) Superposition of domain II of eIF2γ in map A (coloured light blue) with that in map 1 (eIF2γ coloured orange) shows the rearrangement of three eIF2γ domains when it is in the tilted conformation. (f) Conformation of eIF2β fitted in map D (red) is different from the conformations found in three other maps (A-blue, B-green and C-grey). eIF2 γ and α shown are from map D.

To separate the different conformations adopted by eIF2 γ and β, we have applied two masked classifications (Fig. S1 and Methods). After the first masked classification, we obtained three maps A-C (Fig. S1) with conformations of eIF2γ tilted towards the PLD domains of eIF2Bγ subunit and distinct extended conformations of eIF2β (Fig. 3a, 3b and 3f). The tilted conformation of eIF2γ is stabilized by the contacts of eIF2β with the PLD domains of eIF2B γ and ε subunits and the contact of eIF2γ domain III (eIF2γ-D3) with the eIF2Bγ PLD domain (corresponding to residues 97 - 101 and 136 – 139 in eIF2Bγ PLD). This conformation results results in a slight rearrangement of the three domains in eIF2γ, compared to the TC structure (Fig. 3e) ^7,9,41^. Also, the eIF2γ G domain in this conformation is more disordered than in the TC, possibly reflecting a higher mobility of this domain in this particular conformation. Previously a rearrangement of the three γ domains which depended on the nucleotide binding state was reported in a crystallographic study in archaeal aIF2 ^42^.

In all three maps the density for eIF2β allowed modelling of the zinc-binding and central domains in the conformation similar to the one in the TC, but with the zinc-binding domain only partially covering the nucleotide-binding pocket and extended central domain approaching the binding interface between the eIF2B γ and ε PLD domains (Fig. 3f). One of these maps also contained extra density contacting the top of eIF2γ G-domain (Fig. 4a and 4b), large enough to accommodate the ε-cat heat domain in proximity to the N-terminus of eIF2β, previously shown to interact with the ε-cat heat domain ^43^. In this position the ε-cat domain would not have access to the nucleotide-binding pocket on the eIF2γ-G domain. However we cannot exclude the possibility that ε-cat could act allosterically by inducing rearrangement of the domains in eIF2γ, which we can see in the maps with the tilted conformations of eIF2γ, leading to nucleotide release. In this case the 72 residues linker (res. 472–544), connecting ε-cat with the rest of eIF2Bε, is just long enough to cover the distance of around 85 Å that separates this density from the c-terminus of the modelled eIF2Bε (Fig. 4b).

**Figure 4.**
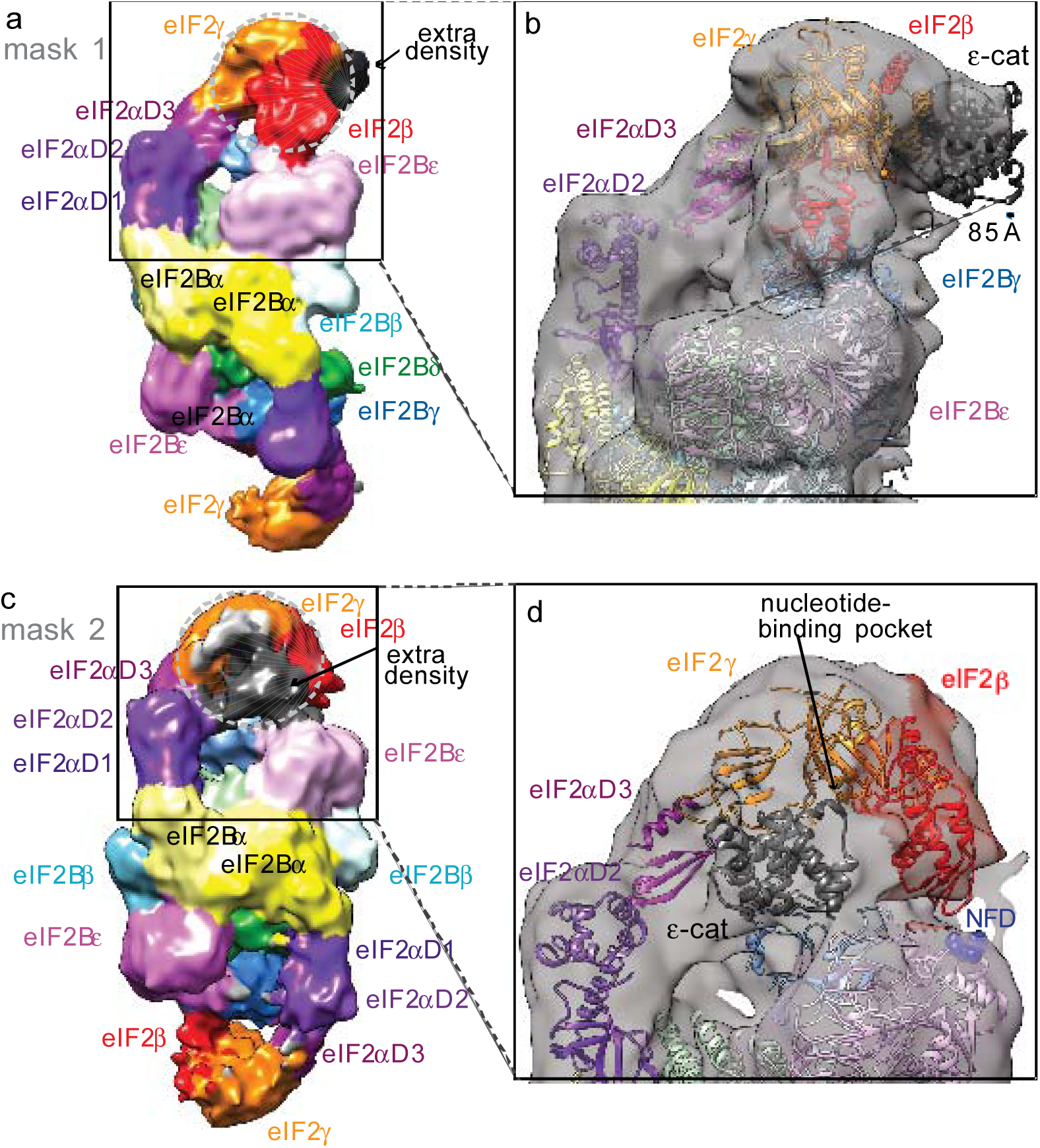
Possible interaction of eIF2B ε-cat heat domain with eIF2. (a) Map B of eIF2B-eIF2(αP) complex obtained by masked classification around eIF2 γ and β showing extra density in contact with eIF2γ. (b) The extra density in map B could accommodate most of the ε-cat heat domain and would be in contact with eIF2γ domains III and G away from nucleotide-binding site. 85 Å distance separates this extra density from the c-terminus of the eIF2Bε and is just enough for the 72 residues linker (res. 472–544) to connect ε-cat with the rest of eIF2Bε. (c) Map D of eIF2B-eIF2(αP) complex obtained by masked classification around eIF2 γ and an extra density seen at a lower threshold (black) in proximity of nucleotide-binding pocket. (d) The size and shape of the extra density in map D will fully account for the whole ε-cat heat domain. Also in this map eIF2β approaches the NFD motif (blue spheres) in eIF2Bε.

A second masked classification yielded a map at only 10.4 Å resolution (Fig. 4c, 4d and S1) but with a quite defined extra density, also of the size of the ε-cat domain, this time, on the other side of the eIF2γ-G domain close enough to the nucleotide binding region (Fig. 4d). Interestingly, this map also contained the density for eIF2β, not included in the mask. In this map eIF2β central domain now approaches the NF motif in eIF2Bε subunit, which is important for catalysis ^34, 36^(Fig. 4d), while zinc-binding domain, although not very well defined, does not cover the nucleotide-binding pocket (red conformation of eIF2β in Fig. 3f).

## Discussion

The structure of eIF2B-eIF2(αP) complex, presented here, directly shows that two eIF2 molecules bind opposite sides of an eIF2B hetero-decamer via spatially separated interactions. Although we do see particles of eIF2B alone, we do not observe particles corresponding to only one molecule of eIF2 bound to eIF2B in our datasets (Fig. S1). This observation suggests that the binding of eIF2 to eIF2B is cooperative. The interaction of eIF2α-D1 to the regulatory moiety of eIF2B is relatively well defined in our structure, and likely makes the major contribution to the affinity between these two factors. In our complex eIF2α was phosphorylated in vitro at Ser51, which is known to result in an even more stable interaction with eIF2B ^12, 40^. The effect of Ser51 phosphorylation may be attributed to a combination of direct interactions with the residues in eIF2Bα and induced contact with eIF2Bδ. The large interaction area of eIF2α-D1 with eIF2B α and δ (844 and 374 Å^2^ respectively), in between which eIF2α-D1 is sandwiched, implies that most of the contacts would be very similar even in the absence of phosphorylation. This conclusion is also supported by cross-linking experiments ^28^ showing that the binding mode of eIF2α to the regulatory moiety of eIF2B is hardly affected by its phosphorylation status. However the additional cross-links which occurred in the absence of phosphorylation to Q91 and R84 of eIF2Bβ identified in the same study ^28^ (corresponding to E100 and A93 in our structure (Fig. 2b) are far from the contact interface, suggesting that the binding of non-phosphorylated eIF2α may not be as stable and possibly leads to a conformational change necessary for nucleotide displacement.

The recently isolated ISR inhibitor (ISRIB) ^44^, was shown by cryoEM to bind human eIF2B at the two-fold symmetric interface “stapling” two βδ dimers of the regulatory core ^26, 27^. ISRIB was shown to boost “catalytic activity” of eIF2B in both phosphorylated and nonphosphorylated eIF2 ^26, 44,45, 46^. Its action was mostly attributed to the stabilization of the eIF2B hetero-decamer in human ^26, 46^, which is less stable than in yeast ^47, 48^. When compared to the eIF2B-eIF2(αP) structure obtained here, it is clear that binding of eIF2 to the eIF2B hetero-decamer, where the binding interfaces for αD1 with eIF2B α, β and δ subunits are preserved, is more stable then in an eIF2B(βδγε) tetramer, where the only binding could be through eIF2Bδ. The higher affinity of decameric eIF2B to eIF2 can explain its higher activity in humans ^48, 49^. Comparison of ISRIB bound eIF2B with our eIF2(αP) bound eIF2B structure (Fig. S3) shows that ISRIB imposes a distinct symmetric eIF2B structure, which is incompatible with stable binding of two eIF2(αP) molecules at the same time (Fig. S 3d and 3e). Rigidifying the eIF2B core by binding ISRIB across two eIF2B βδ dimers may also preclude complete closure of eIF2Bδ helical bundle CTD around eIF2α-D1 reducing the contact area and thereby the affinity to eIF2α, independent from its phosphorylation status at least in one of the eIF2 molecules.

Previously eIF2(αP) has been shown to effectively sequester eIF2B ^50, 51^, but also act as a competitive inhibitor of nucleotide exchange and prevent catalysis by non-productive interactions of eIF2(αP) with eIF2Bε-cat ^21^. The local resolution in eIF2 γ, β and eIF2B ε-cat does not allow us to elucidate the details of nucleotide displacement. However, inhibition of the nucleotide exchange by eIF2α phosphorylation in the same molecule would not account on its own, for relatively small proportion of phosphorylated eIF2 (~30 %) sufficient for inhibiting eIF2B activity ^51^, as the majority of non-phosphorylated eIF2 still would be available for productive nucleotide exchange even with limiting amounts of eIF2B in the cell. In contrast, the idea of sequestration of the much less abundant eIF2B when compared to eIF2 (10 times less ^52^) in a non-productive complex, seems the most important reason for translation inhibition by eIF2 phosphorylation, especially since binding of eIF2(αP) to the regulatory subunits of eIF2B is enhanced when compared to its unphosphorylated form and necessary for the inhibition of translation ^12^.

Recently Jennings et al showed that nucleotides have a minor impact on the overall affinity of eIF2 to eIF2B ^53^, likely reflecting the fact that binding of eIF2 to the regulatory core of eIF2B through α-D1 makes the major contribution to the affinity and not the interactions with the catalytic eIF2B subunits. Our reconstructions of the eIF2B-eIF2(αP) complex show high mobility and flexibility of eIF2 γ and β around catalytic portion of eIF2B, while maintaining the stronger contact through eIF2α-D1. In the cell the probability of GTP binding by eIF2 after GDP displacement by the catalytic portion of eIF2B is much higher than that of GDP due to an approximately 10 times higher GTP concentration. This would allow Met-tRNA_i_^Met^ binding to eIF2γ while eIF2α-D1 is still being attached to regulatory portion of eIF2B. Indeed, thes acceptor stem of Met-tRNA_i_^Met^ mainly contributes to the affinity of eIF2 binding in the TC ^54, 55^, suggesting that this contact is driving formation of the TC. Superposition of eIF2 bound to eIF2B in our complex with the eIF2 structure in the TC ^7^ show that this interaction is possible in the context of eIF2B-eIF2(αP) complex (Fig. 5a). In fact a stable interaction of eIF2B in complex with GTP-eIF2 and Met-tRNA_i_^Met^ has been shown previously ^50^.

**Figure 5.**
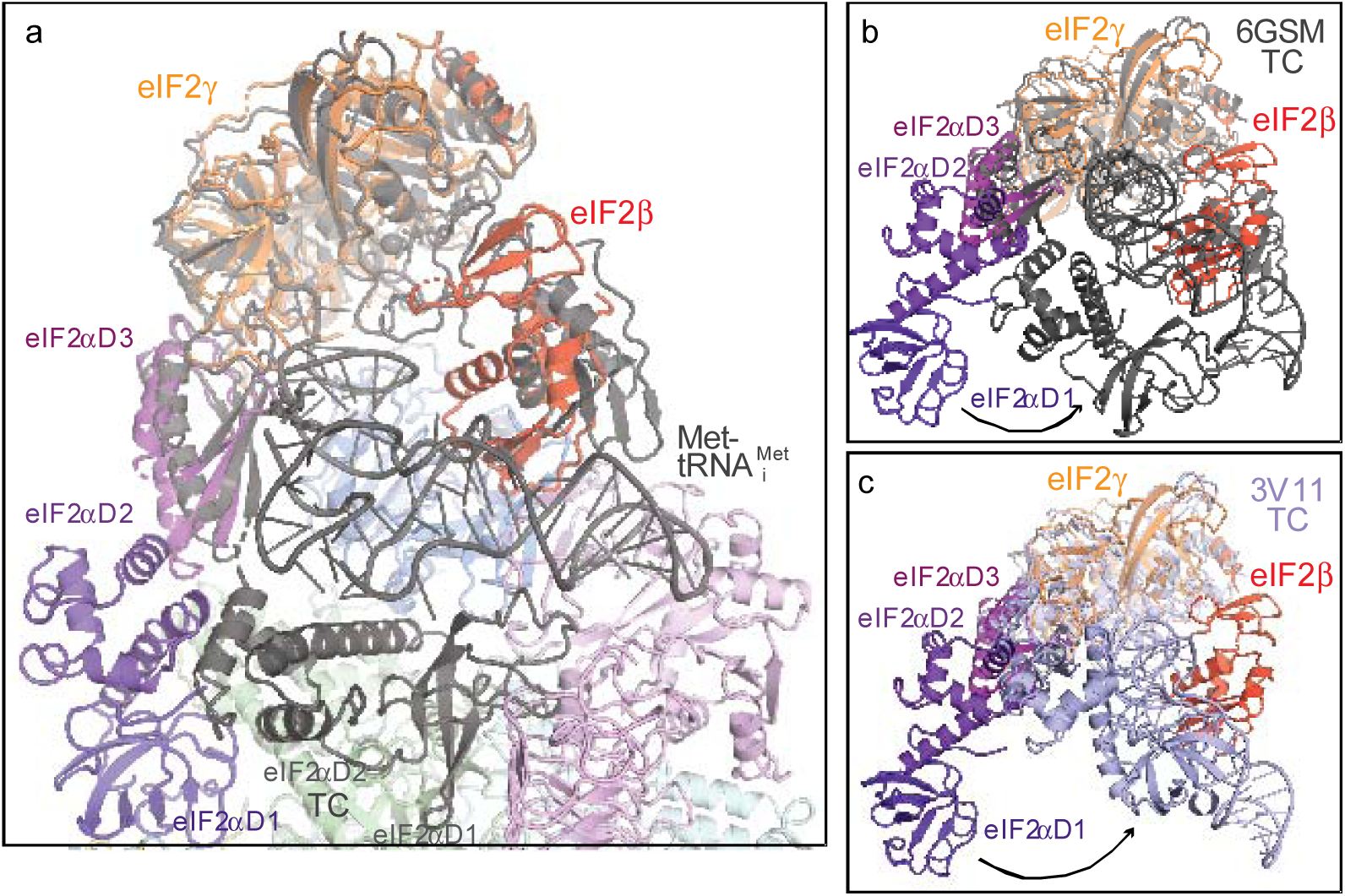
Superposition of the TC with eIF2B - eIF2(αP) complex based on eIF2γ. (a) Superposition of yeast TC (6GSM) in grey with the model of eIF2B - eIF2(αP) complex in map C showing that Met-tRNA_i_^Met^ can bind without clash to eIF2γ and eIF2α- D3 while eIF2α-D1 is still attached to eIF2B. (b) Same superposition as in (a) in a different orientation shows the large conformational changes that eIF2α - D1 and D2 undergo when bound to eIF2B or Met-tRNA_i_^Met^ both competing for eIF2α. (c) Same as in (b), but superimposed with *S. solfataricus* TC (3V11, light blue).

For the completion of the TC formation, a large conformational change in eIF2α is needed (Fig. 5b and 5c), since eIF2α-D1 must be extracted from eIF2B, as both Met-tRNA_i_^Met^ and eIF2Bδ share the same binding interface with the helix 58–63 in eIF2α. This shared binding interface creates direct competition between Met-tRNA_i_^Met^ and eIF2B for eIF2 binding. The competition between eIF2B and Met-tRNA_i_^Met^ for eIF2 binding has been recently shown experimentally ^53^. Our structure suggests that Ser51-P directly interacts with residues in eIF2Bα, and that phosphorylation of eIF2α Ser51 slightly alters the conformation of the helix 58–63 in eIF2α, which may tip the balance towards eIF2B binding and prevent TC formation.

At the same time the competition for eIF2 between eIF2B and Met-tRNA_i_^Met^ is also influenced by the competition for eIF2 between eIF2B ε-cat and eIF5-CTD ^56, 43, 57^, which share the same fold. Both eIF2B ε-cat and eIF5-CTD bind the eIF2γ-G domain as well as the same region in eIF2β ^56, 43, 57^, the former displacing the nucleotide and the latter protecting it from displacement ^58, 53,59^. While eIF2B was shown to disrupt TC ^53^, however adding eIF5 or eIF5-CTD to the TC protected it from disruption, but not when eIF2α is phosphorylated. These data suggest that there is a fine balance between the catalytic and regulatory interactions of eIF2 and eIF2B, which are affected by other binding partners – eIF5 and Met-tRNA_i_^Met^. We propose, that it is not eIF2B that discriminates between the nucleotide states of eIF2, but rather subsequent interactions with Met-tRNA_i_^Met^ allow this discrimination in the cell. In fact the presence of Met-tRNA_i_^Met^ has been shown to stimulate the rate of GDP to GTP exchange by eIF2B ^50, 60^.

Sequestering of eIF2B by phosphorylated eIF2, which is present in cell in ~ 10 times excess, has been suggested as a mechanism of ISR based on a number of biochemical studies ^12, 51^ and generally is in agreement with the structure of eIF2B-eIF2(αP) complex that we have obtained. However, the sequestration does not necessarily have to be irreversible. A slow dissociation rate of eIF2(αP) would prevent high turnover of eIF2B recycling and subsequent binding to non-phosphorylated eIF2. Therefore, the picture emerges that GEF and ISR function of eIF2B are structurally coupled and driven kinetically by the further formation of the TC (and eIF5 binding) – proceeding to initiation.

## Acknowledgments

We thank G. Pavitt for providing *Saccharomyces cerevisiae* strain GP4109 for over-expressing yeast eIF2B. We are grateful to G. Cannone and G. McMullan for technical support with cryo-EM, T. Darling and J. Grimmett for help with computing.

## Author contributions

YG purified the protein complex and prepared the samples. JLL and YG performed electron cryo-microscopy data collection. JLL performed processing, model building and analysis. YG and JLL wrote the first draft of the manuscript. VR provided support for the research and helped edit and revise the manuscript.

## Data availability

Six maps have been deposited in the EMDB with accession codes EMDB: 1111, EMDB: 2222, EMDB: 3333, EMDB: 4444, EMDB: 5555, EMDB: 6666, for Map 1, Map 2, Map A, Map B, Map C and Map D, respectively. Five atomic coordinate models have been deposited in the PDB with accession codes PDB: XXXX, PDB: YYYY, PDB: ZZZZ, PDB: AAAA, PDB: BBBB for Maps 1 & 2, Map A, Map B, Map C and Map D, respectively.

## Supplementary Materials

Figures S1-S5

## Methods

### Protein purification and complex assembly

*Saccharomyces cerevisiae* eIF2 was purified from yeast strain GP3511 (*MAT*α *leu2-3 leu2-112 ura3-52::HIS4-lacZ ino1 gcn2*Δ *pep4::LEU2 sui2*Δ pAV1089[*SUI2 SUI3 GCD11*-His6 2 μm *URA3*] ^1^ as described previously ^2^. Prior to assembly of the complex with eIF2B, purified eIF2 was phosphorylated *in vitro* by human PKR (Invitrogen) ^3^.

*Saccharomyces cerevisiae* eIF2B was over-expressed in yeast strain GP4109 (*MAT*α *leu2-3 leu2-112 ura3-52 ino1 gcd6*Δ *gcn2*Δ::hisG *ura3-52::HIS4*-*lacZ* pAV1428[*GCD6 GCD1*-FLAG2-His6 *URA3* 2 μm] pAV1494[*GCN3 GCD2 GCD7 LEU2* 2 μm]) ^4^. After harvesting cells were suspended 1:1 (w:v) in PBS and cell suspension droplets were frozen in liquid nitrogen. Usually 50g of cell “popcorn” was used for each purification of the eIF2B in complex with phosphorylated eIF2 (eIF2(αP)). After cell lysis Flag-tagged eIF2B complexes were immobilised on 300 μl of Anti-Flag M2 affinity gel (Sigma) and washed with a high-salt buffer (500 mM KCl) ^5^ followed by phosphorylation buffer (20 mM Tris (pH 7.5), 100 mM KCl, 10 mM MgCl2, 5 mM β-glycerophosphate, 2 mM dithiothreitol, 10% glycerine, 0.1% NP-40, 200 μM ATP). The amount of eIF2B was estimated not to exceed 200 μg from 50g of cell “popcorn” based on repeated purifications of eIF2B on its own. Therefore in our phosphorylation reaction we used over ~ 2 times equimolar amount of eIF2 assuming two molecules of eIF2 can bind one eIF2B hetero-decamer. Phosphorylation buffer containing 2 mM GDP was added to immobilised Flag-tagged eIF2B and incubated at room temperature for 20 min. Beads were washed two times with the buffer - 20 mM Hepes (pH7.5), 100 mM KCl, 5mM MgCl_2_, 5mM β-ME. eIF2B-eIF2(αP) complexes were eluted in 250 μl of the same buffer containing 100 μg/ml of 3XFlag peptide (Sigma) and washed/concentrated in Amicon Ultra 50K MWCO concentrators 5 times in the buffer without 3XFlag-peptide. Protein concentration was measured by nanodrop and Bradford reaction, which gave concentration values within 10% difference, usually in the range of 1 to 2 μg/μl or 1.17 to 2.35μM (assuming two molecules of eIF2 bind eIF2B hetero-decamer) (Fig. S5a).

Immediately before applying to cryo grids, the sample was diluted to ~ 200 nM with the same buffer containing glutaraldehyde (~ 0.125%) to make the final concentration of glutaraldehyde 0.1 %.

### Electron microscopy

Three μl of the eIF2B-eIF2(αP) complex were applied to glow-discharged gold UltrAuFoil R 1.2/1.3 or R 2/2 grids at 4 °C and 100% ambient humidity. After 30s incubation, the grids were blotted for 4-5 s and vitrified in liquid ethane using a Vitrobot Mk3 (FEI).

Automated data acquisition was done using the EPU software (FEI) on a Titan Krios microscope (FEI) operated at 300 kV under low-dose conditions in linear (data set I, 45 e^−^/Å^2^) or counting mode (data set II, 21 e^−^/Å^2^) using a defocus range of 1.5 – 4.5 μm. In linear mode, images of 1.1 s/exposure and 34 movie frames were recorded (Fig. S5b), whereas in counting mode, we saved 75 fractions over a 60 second exposure, using in both cases a Falcon III direct electron detector (FEI) at a calibrated magnification of 104,478 (yielding a pixel size of 1.34 Å). Micrographs that showed noticeable signs of astigmatism or drift were discarded.

### Analysis and structure determination

The movie frames were aligned with MotionCor2 ^6^ for whole-image motion correction. Contrast transfer function parameters for the micrographs were estimated using Gctf ^7^. Particles were picked using Relion ^8^. References for template-based particle picking ^9^ were obtained from 2D class averages that were calculated from particles semi-automatically picked with EMAN2 ^10^ from a subset of the micrographs. 2D class averaging (Fig. S5c), 3D classification and refinements were done using RELION-2 ^8^. Both movie processing ^11^ in RELION-2 and particle ‘‘polishing’’ were performed for all selected particles for 3D refinement. Resolutions reported here (Fig. S5d) are based on the gold-standard FSC = 0.143 criterion ^12^. All maps were further processed for the modulation transfer function of the detector, and sharpened ^13^. Local resolution was estimated using ResMap ^14^.

For the data set I, 3,282 images were recorded from two independent data acquisition sessions, and 459,480 particles were selected after two-dimensional classification. An initial 3D reconstruction was made from all selected particles after 2D class averaging using the *Schizosaccharomyces pombe* eIF2 cryoEM structure (PDB: 5B04) low-pass filtered to 60Å as an initial model, and using internal C2 symmetry. Next, two consecutive 3D classification into 15 and 6 classes, respectively, this time without using the eIF2B internal symmetry, with a 7.5 degrees angular sampling interval and no local searches was performed to remove bad particles or empty eIF2B particles from the data and to get an initial understanding of the conformational heterogeneity of eIF2 in the complex. After the second round of 3D classification 239,695 particles were selected (52% of the total) and refined to 5.7 Å-resolution

The map did not yield a high overall resolution, so we decided to collect an additional dataset at the same magnification and using the same detector but in counting instead of linear mode. For this dataset (data set II), 1,241 images were recorded, and 173,740 particles were selected after two-dimensional classification. After obtaining an initial three-dimensional refined model, and two consecutive rounds of 3D classification the classes containing the eIF2B-eIF2(αP) complex were selected (131,663 particles, 75 % of the total) and after movie processing, refined using C2 internal symmetry to much higher resolution than for the data set I (Map I, 4.15 Å).

The particles from both datasets were then combined and a masked 3D classification using masks around two eIF2γ molecules in the complex was carried out to remove particles with low occupancy for these factors, as a result of which 183,468 particles were selected and refined to 4.25 Å (Map II). The overall resolution of this map was slightly lower than that of map I, but the occupancy and local resolution for eIF2γ and eIF2α-D3 was better.

The preliminary 3D rounds of classification showed that eIF2γ, eIF2α-D3, and densities possibly belonging to eIF2β and the heat domain of eIF2Bε adopt many different conformations. So we carried out 3D classifications with subtraction of the residual signal ^15^ by creating two different masks around the density attributed to eIF2α-D3, eIF2γ and eIF2β in all possible conformations observed in the preliminary 3D classification rounds, and around a density observed at low threshold in close proximity to the eIF2γ G-domain. We applied these masks for each of the two molecules of eIF2 in each eIF2B-eIF2(αP) complex. We isolated four distinct and well-defined maps by ‘focused’ 3D classifications, as follows.

A. Map A, showing higher occupancy for eIF2β and a tilted conformation of eIF2γ [119,037 particles, 4.55Å],
B. Map B, similar to map A but with slightly different conformations of eIF2β and eIF2γ. It also shows an extra density in contact with the G-domain and domain III of eIF2γ [12,575 particles, 9.4Å].
C. Map C, showing the most extreme tilted conformation towards eIF2Bγ for eIF2γ, and where eIF2β is also observed [23,909 particles, 10.1Å], and,
D. Map D, showing additional density in contact with eIF2γ, whose size and shape suggested that it could correspond to eIF2B ε-cat heat domain [23,909 particles, 10.4Å]

### Model building and refinement

In all six maps the conformations of all eIF2B subunits and domains D1 and D2 of eIF2α are nearly identical. Thus, modelling of all these elements was first done in the higher resolution maps (4.15 and 4.25 Å; Maps 1 and 2), and then this model was used as a reference for model building in EM maps with lower resolution (maps A to D). In this procedure, the cryoEM model of eIF2B from *S. pombe* (PDB: 5B04) was placed into density by rigid-body fitting using Chimera ^16^. Then each subunit of eIF2B was independently fitted by rigid-body refinement, first in Chimera and then in Coot ^17^. Also in Coot, the sequence was converted to that of *S. cerevisiae* proteins, followed by rigid body of different subdomains within each eIF2B subunit. Further modelling was also done in Coot, paying special attention to the region of eIF2B in contact with eIF2.

eIF2 was taken from PDB: 6FYX. eIF2α-D1/eIF2α-D2 and eIF2α-D3/eIF2γ/eIF2β N-terminal helix were fitted as separate rigid bodies into its corresponding densities, using Chimera and Coot. Then, each of these domains but the eIF2β n-terminal helix was independently fitted, and further modelling was also done in Coot.

Model refinement in the highest resolution maps was carried out in Refmac v5.8 optimized for electron microscopy ^18^, using external restraints generated by ProSMART ^18^. The average Fourier Shell Coefficient (FSC) was monitored during refinement. The final model was validated using MolProbity ^19^. Cross-validation against overfitting was done as previously described ^18, 20^. Refinement statistics for the last refinements, done in Map 2, are given in Table 1.

**Table 1.** Refinement and model statistics.

|  | **Model refined in**<br>**maps 1 and 2** | **Model refined in**<br>**map A** |
| --- | --- | --- |
| **Model Composition** |  |  |
| Non-hydrogen atoms | 37,066 | 37,959 |
| Protein residues | 4,964 | 5,075 |
| **Refinement** |  |  |
| Resolution used for refinement (Å) | 4.25 | 4.60 |
| Map sharpening B-factor (Å) | -119 | -100 |
| Average B-factor (Å) | - | - |
| Fourier Shell Correlation (FSC)* | 0.45 | 0.42 |
| **Rms deviations** |  |  |
| Bonds (Å) | 0.008 | 0.008 |
| Angles (°) | 1.215 | 1.258 |
| **Validation (proteins)** |  |  |
| Molprobity score<br>(Percentile in brackets) | 2.33<br>(99<sup>th</sup>) | 2.50<br>(98<sup>th</sup>) |
| Clashscore, all atoms<br>(Percentile in brackets) | 3.09<br>(100<sup>th</sup>) | 3.68<br>(100<sup>h</sup>) |
| Good rotamers (%) | 81.1 | 78.8 |
| **Ramachandran plot** |  |  |
| Favored (%) | 87.0 | 83.3 |
| Outliers (%) | 2.9 | 3.5 |

\* $FSC = \Sigma(N_{shell} FSC_{shell}) / \Sigma(N_{shell})$ , where $FSC_{shell}$ is the FSC in a given shell, $N_{shell}$ is the number of “structure factors” in the shell. $FSC_{shell} = \Sigma(F_{model} F_{EM}) / (\sqrt{\Sigma(|F|_{model}^2)} \sqrt{\Sigma(|F|_{EM}^2)})$

These refined models were used as initial models for maps A-D, and then each subunit of the model was rigid body fitted, without observing almost any appreciable change, except for the eIF2α-D3/eIF2γ/eIF2β n-terminal helix sub-module in one of the two eIF2 molecules. After the fitting of this eIF2α-D3/eIF2γ/eIF2β n-terminal helix sub-module in each of these maps, an extra density belonging to the whole eIF2β subunit was observed and we consequently docked into it the subunit β from PDB: 6FYX. In map D, although there is density for most of eIF2β, it was not possible to do an appropriate rigid body docking without any major clashes and we decided not to include eIF2β in the final model. We also decided not to include eIF2B ε-cat heat domain in any of the models in maps B or D due to its poor local resolution. All figures were generated using PyMOL, Coot or Chimera.

