## Supplemental Figures S1 to S5 for "Structural basis for the inhibition of translation through eIF2α phosphorylation"

**Figure S1**. 3D- classification scheme.

**Figure S2**. Local resolutions of eIF2B - eIF2(αP) complex.

(a) Local resolutions of eIF2B - eIF2(αP) complex in map 2 (related to Fig. 1a) showing surface (left) and two cross-sections around eIF2α-D1 binding site and location of Ser-51.

(b) Same as in (a), but in in map A (related to Fig. 1b).

**Figure S3.** Superposition of the ISRIB bound human eIF2B to eIF2B - eIF2(αP) complex.

(a) Superposition of eIF2B-eIF2(αP) complex with human ISRIB bound eIF2B (black) (PDB 6CAJ) showing elongation of eIF2B hetero-decamer towards catalytic poles upon binding of eIF2 by ~ 12 Å.

(b) Same superposition as in (a), but also including *S. pombe* eIF2B structure (grey) (PDB 5B04), shows that elongation of eIF2B is mainly induced by closure eIF2B α and δ around eIF2α-D1 displacing eIF2Bγ outwards.

(c) Same superposition as in (a), showing displacement of eIF2Bγ in eIF2B-eIF2(αP) complex in one of the eIF2B poles.

(d) The binding site of eIF2α-D1 contacting the superposed eIF2Bα subunit showing reduced contacts with eIF2Bδ and local rearrangement of eIF2Bα interacting helices (indicated with an arrow).

(e) The other binding site of eIF2α-D1 showing that on this side the binding pocket for eIF2α-D1 formed by eIF2B α and δ is wide open.

**Figure S4**. Examples of structural models fitting in density maps.

(a) Density for the contact of eIF2α-D1 with eIF2B α and δ subunits.

(b) Density for Ser51(P).

(c) Density for eIF2Bε PLD domain.

(d) Density for eIF2Bε LβH domain.

(e) Density for eIF2Bδ in contact with eIF2Bβ and eIF2Bα subunits.

(f) Density for eIF2Bα subunit.

**Figure S5**. Methods.

(a) SDS PAGE of eIF2B - eIF2(αP) complex with eIF2B subunits labelled in black and eIF2 subunits in red.

(b) Typical micrograph of eIF2B - eIF2(αP) complex (see methods for details).

(c) 2-D classification of eIF2B - eIF2(αP) complex.

(d) FSC curves.

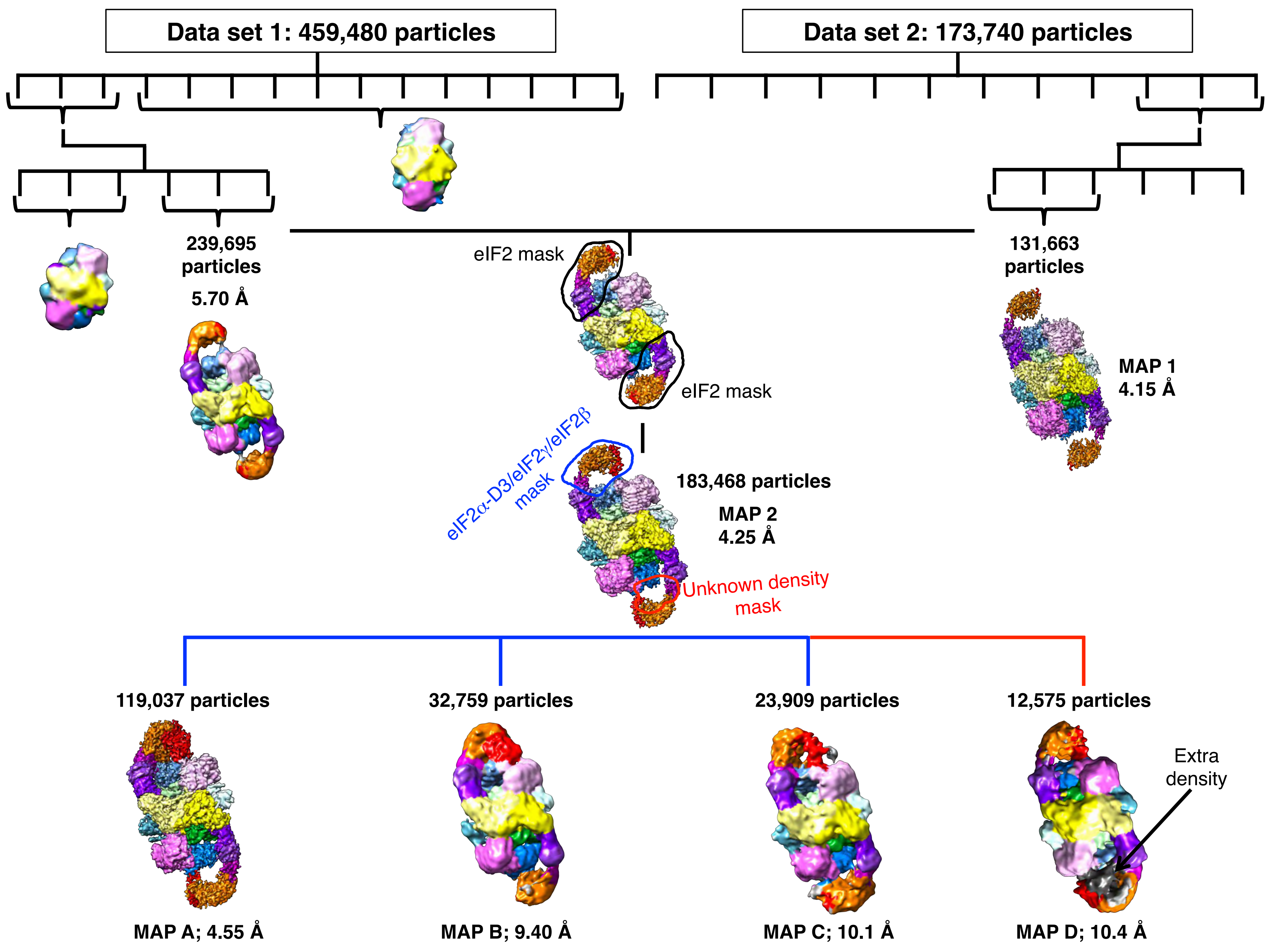

Data set 1: 459,480 particles
Data set 2: 173,740 particles
239,695 particles
5.70 Å
eIF2 mask
eIF2 mask
131,663 particles
MAP 1 4.15 Å
eIF2α-D3/eIF2γ/eIF2β mask
183,468 particles
MAP 2 4.25 Å
Unknown density mask
119,037 particles
32,759 particles
23,909 particles
12,575 particles
Extra density
MAP A; 4.55 Å
MAP B; 9.40 Å
MAP C; 10.1 Å
MAP D; 10.4 Å

Figure S2

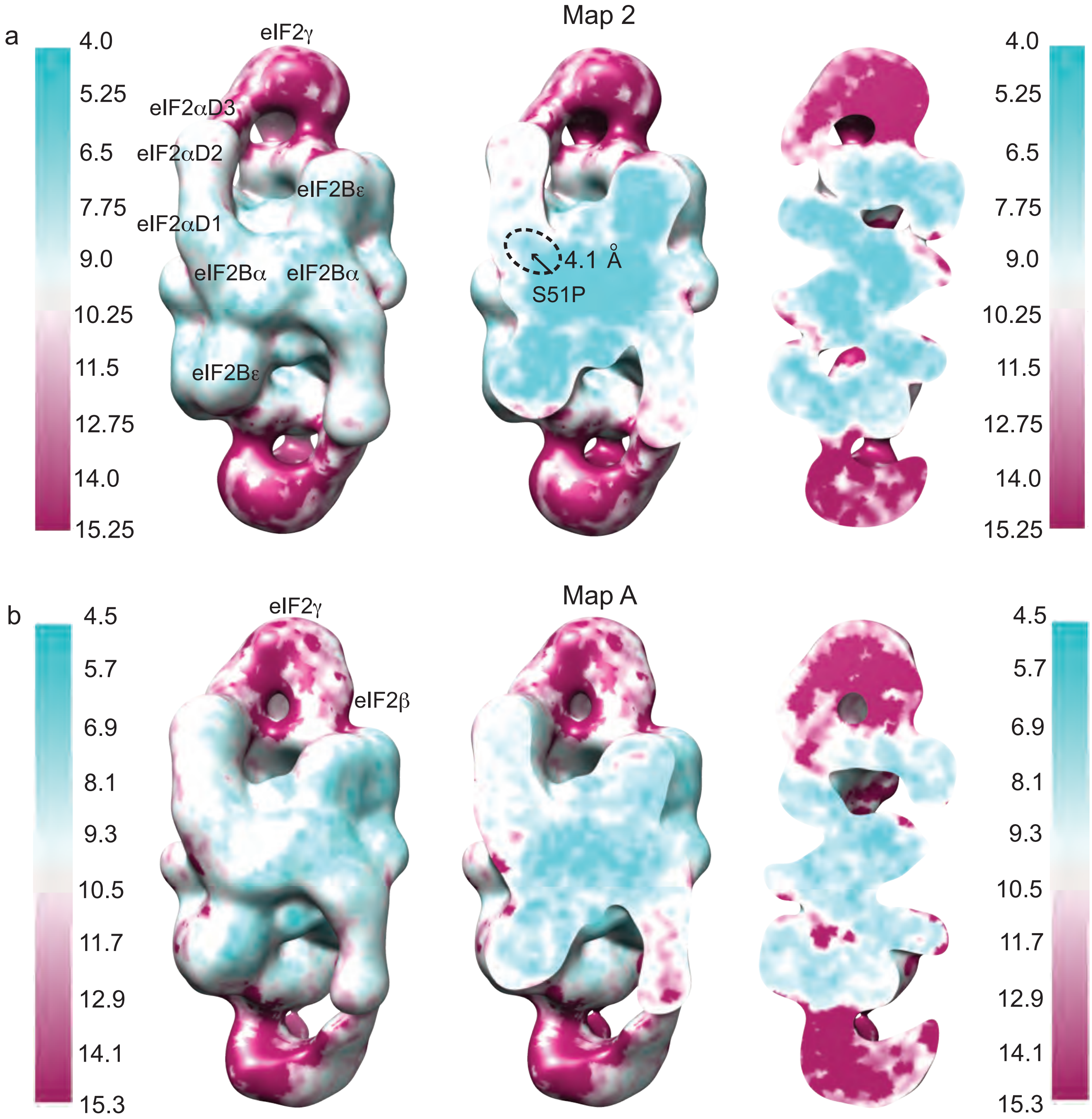

a
4.0
5.25
6.5
7.75
9.0
10.25
11.5
12.75
14.0
15.25
eIF2γ
eIF2αD3
eIF2αD2
eIF2Bε
eIF2αD1
eIF2Bα
eIF2Bα
eIF2Bε
Map 2
4.1 Å
S51P
b
4.5
5.7
6.9
8.1
9.3
10.5
11.7
12.9
14.1
15.3
eIF2γ
eIF2β
Map A

Figure S3

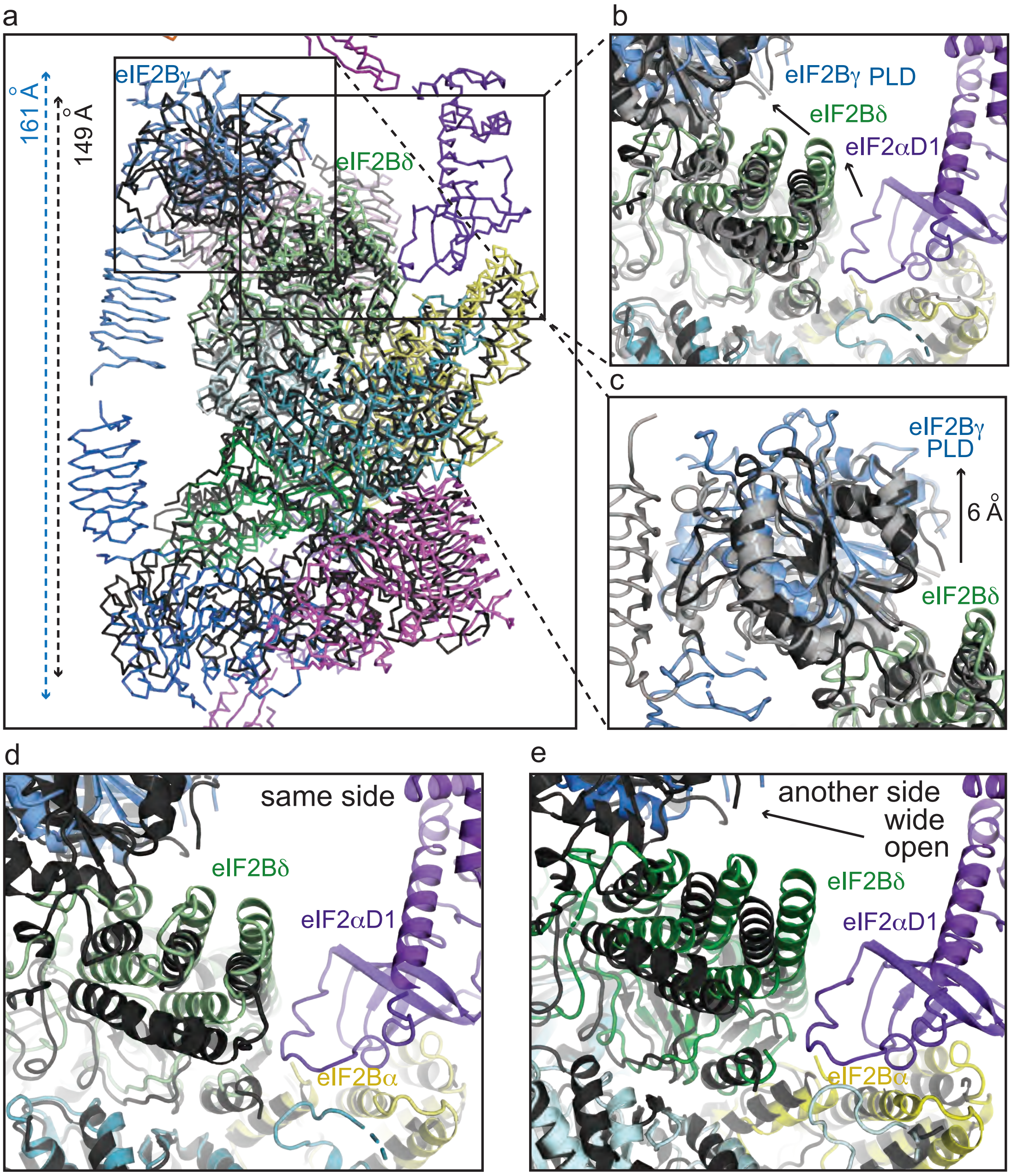
a
b
c
d
e
eIF2Bγ
161 Å
149 Å
eIF2Bδ
eIF2Bγ PLD
eIF2Bδ
eIF2αD1
eIF2Bγ
PLD
6 Å
eIF2Bδ
same side
eIF2Bδ
eIF2αD1
eIF2Bα
another side
wide
open
eIF2Bδ
eIF2αD1
eIF2Bα

Figure S4

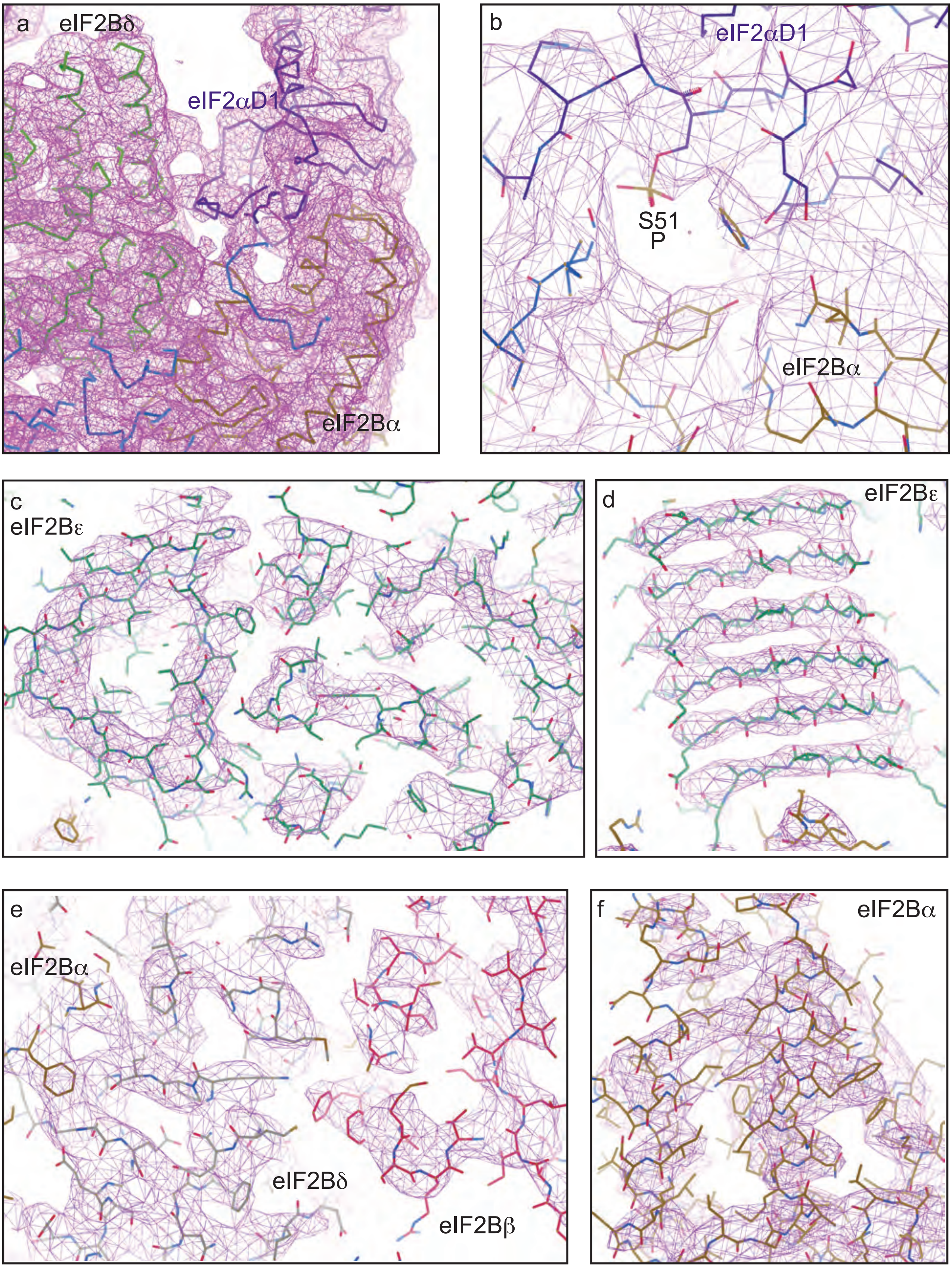
a
eIF2Bδ
eIF2αD1
eIF2Bα
b
eIF2αD1
S51
P
eIF2Bα
c
eIF2Bε
d
eIF2Bε
e
eIF2Bα
eIF2Bδ
eIF2Bβ
f
eIF2Bα

Figure S5

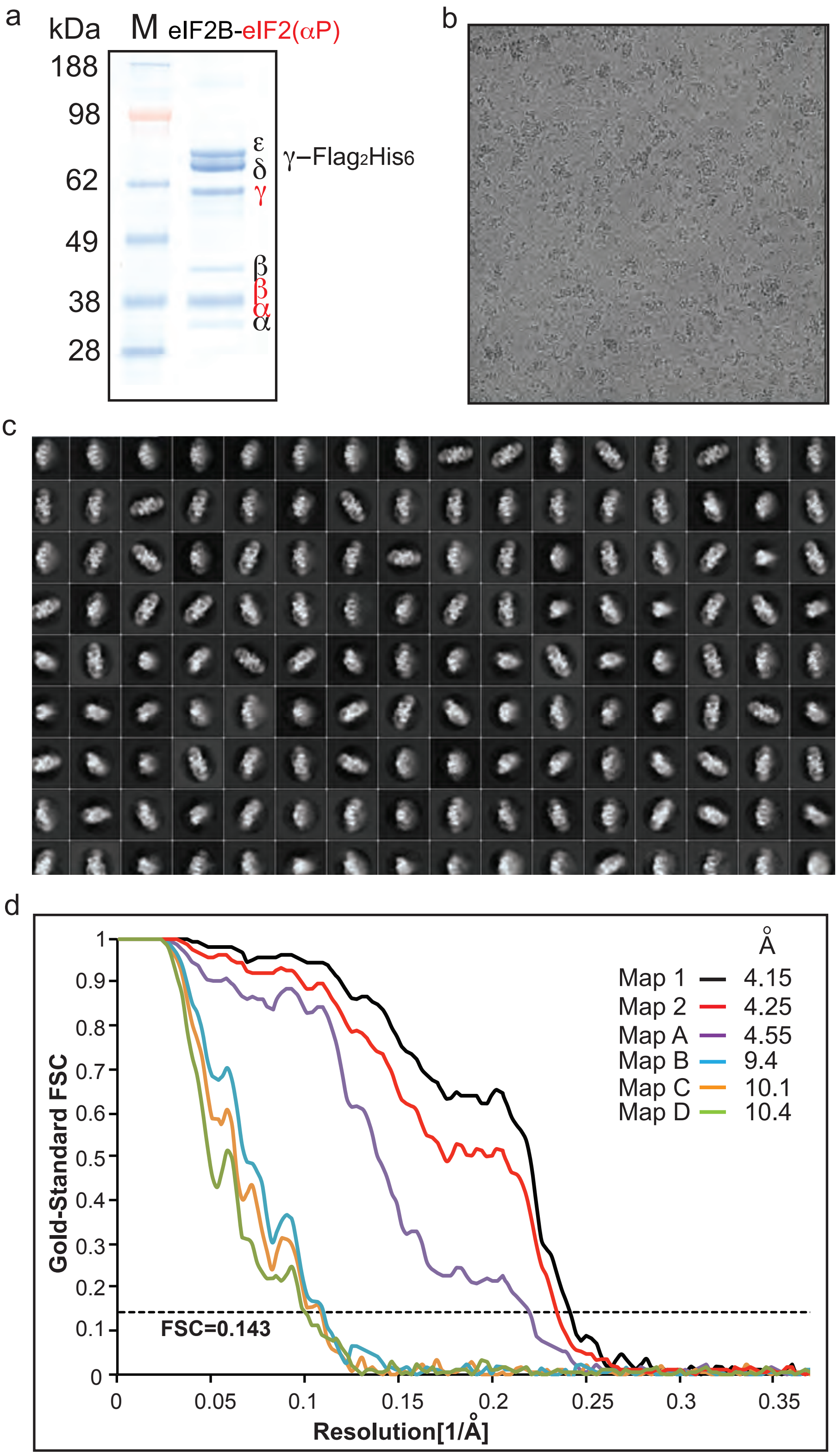

a
kDa
M
eIF2B-eIF2(αP)
188
98
62
49
38
28
ε
δ
γ
β
β
α
α
γ−Flag2His6
b
c
d
1
0.9
0.8
0.7
0.6
0.5
0.4
0.3
0.2
0.1
0
Gold-Standard FSC
FSC=0.143
0
0.05
0.1
0.15
0.2
0.25
0.3
0.35
Resolution[1/Å]
Å
Map 1 4.15
Map 2 4.25
Map A 4.55
Map B 9.4
Map C 10.1
Map D 10.4
